# A Genome Sequence for the Threatened Whitebark Pine

**DOI:** 10.1101/2023.11.16.567420

**Authors:** David B. Neale, Aleksey V. Zimin, Amy Meltzer, Akriti Bhattarai, Maurice Amee, Laura Figueroa Corona, Brian J. Allen, Daniela Puiu, Jessica Wright, Amanda R. De La Torre, Patrick E. McGuire, Winston Timp, Steven L. Salzberg, Jill L. Wegrzyn

## Abstract

Whitebark pine (WBP, *Pinus albicaulis*) is a white pine of subalpine regions in western contiguous US and Canada. WBP has become critically threatened throughout a significant part of its natural range due to mortality from the introduced fungal pathogen white pine blister rust (WPBR, *Cronartium ribicola*) and additional threats from mountain pine beetle (*Dendroctonus ponderosae*), wildfire, and maladaptation due to changing climate. Vast acreages of WBP have suffered nearly complete mortality. Genomic technologies can contribute to a faster, more cost-effective approach to the traditional practices of identifying disease-resistant, climate-adapted seed sources for restoration. With deep-coverage Illumina short-reads of haploid megametophyte tissue and Oxford Nanopore long-reads of diploid needle tissue, followed by a hybrid, multistep assembly approach, we produced a final assembly containing 27.6 Gbp of sequence in 92,740 contigs (N50 537,007 bp) and 34,716 scaffolds (N50 2.0 Gbp). Approximately 87.2% (24.0 Gbp) of total sequence was placed on the twelve WBP chromosomes. Annotation yielded 25,362 protein-coding genes, and over 77% of the genome was characterized as repeats. WBP has demonstrated the greatest variation in resistance to WPBR among the North American white pines. Candidate genes for quantitative resistance include disease resistance genes known as nucleotide-binding leucine-rich-repeat receptors (NLRs). A combination of protein domain alignments and direct genome scanning was employed to fully describe the three subclasses of NLRs (TNL, CNL, RNL). Our high-quality reference sequence and annotation provide a marked improvement in NLR identification compared to previous assessments that leveraged de novo assembled transcriptomes.

## Introduction

Whitebark pine (*Pinus albicaulis*) is a five-needle pine of subgenus *Strobus*, section *Quinquefoliae*, subsection *Strobus*. Sugar pine (*Pinus lambertiana*) is a closely related member of the same subsection whose genome was previously sequenced (Stevens et al. 2016). Whitebark pine is found in subalpine regions in western contiguous US and Canada and is most often the tree-line tree species where it occurs. Whitebark pine is of significant and somewhat unique ecological importance. Its wingless seeds are harvested, dispersed, and cached by the Clark’s nutcracker (*Nucifraga columbiana*). Thus, there is a mutualism between the tree and the bird to the extent they have coevolved (Tomback et al. 2001). In areas of joint whitebark pine, red squirrel (*Tamiasciurus hudsonicus*) and grizzly bear (*Ursus arctos horribilis*) habitat, whitebark pine seeds from cones cached in squirrel middens are an important food source for the bears (Mattson and Reinhart 1997). In addition, whitebark pine trees provide shade to the winter snowpack that helps extend the length of the annual snowmelt.

Unfortunately, for all its great ecological importance to the subalpine environment, whitebark pine has become critically threatened throughout a significant part of its natural range (Tomback et al. 2001). The primary threat is mortality due to the introduced fungal pathogen white pine blister rust (*Cronartium ribicola*). Additional threats include mountain pine beetle (*Dendroctonus ponderosae*), wildfire, and maladaptation due to changing climate (Tomback and Achuff 2010). At some locations in the Northern Rockies and Canada, vast acreages of whitebark pine have suffered nearly complete mortality. In December 2022, after years of conservation efforts by the Whitebark Pine Ecosystem Foundation (whitebarkfound.org) and American Forests (americanforests.org), the United States Fish and Wildlife Service listed whitebark pine as a threatened species (US FWS 2022).

There is now an urgent need to conserve and restore whitebark pine throughout its natural range. This can be effectively accomplished if a very large number of white pine blister rust resistant and climate adapted seed sources can be identified, and if planting stock can be produced from those sources. Forest resource managers have for many years been developing such resources using phenotypically-based approaches. Identifying white pine blister rust resistant sources involves finding putatively resistant trees in natural stands, collecting seed from those trees, producing seedlings and artificially inoculating seedlings with blister rust (Sniezko et al. 2008). This approach has been effective in several white pine species, notably sugar pine and western white pine (*Pinus monticola*), however, the discovery process might take 5 to 10 years to complete with a cost of ∼$1500/tree. Likewise, identifying climate-adapted sources employs long-term genetic testing in common gardens that can take decades to complete (Bower and Aitken 2008). Thus, any new technology that could speed up and reduce the cost of identifying seed sources for restoration would be highly desired. Genomic technologies offer one such solution. Just as has been done for human disease screening and for agronomically important traits in domestic crops and livestock, the specific genes underlying these traits must first be discovered. This is the long-term goal for our research. However, this discovery is profoundly enhanced by having a well assembled and annotated reference genome sequence. To that end, in this paper we report on the first reference genome sequence for whitebark pine.

## Materials and Methods

### Reference tree

An approximately 150-year-old tree was selected from the Deschutes National Forest near Bend, Oregon by a USDA Forest Service geneticist. The exact identification number and location of the tree is held in confidence to maintain its security. Scion from the tree was collected and grafted to rootstock; clones are maintained at the USDA Forest Service Dorena Genetic Resource Center in Cottage Grove, Oregon. Tissue from these clones can be obtained upon request. Cones and needle tissue were collected from the reference tree in 2006 and 2021, respectively.

### DNA isolation

The protocol used to isolate the haploid megagametophyte tissue from a single fertilized whitebark pine seed was similar to previous conifer genome sequencing projects (Neale et al. 2014; Zimin et al. 2014). Haploid genomic DNA was extracted from a single megagametophyte with the Omega Biotek E.Z.N.A.®SP Plant DNA Kit. The extraction followed the manufacturer’s protocol with the following modifications: polyvinylpyrrolidone (0.01g) was added to the tissue prior to lysis, and the lysis time was extended to 1.5 hours. The extracted DNA was quantified on a Qubit 2.0 (42.2 ng/μl), a Nanodrop ND-1000 (A260/280: 1.83; A260/230: 2.11), and quality was evaluated on an electrophoresis gel (fragment sizes >20,000 bp).

### DNA sequencing

#### Illumina short read

DNA was sequenced at the DNA Technologies and Expression Analysis Core at the UC Davis Genome Center. First, DNA libraries were made that were suitable for whole genome shotgun sequencing with no unique molecular identifiers. Then sequencing was conducted on 3.5 lanes of a NovaSeqS4 with Illumina 150 bp paired ends sequencing with an approximate insertion size of 400 bp, nonoverlapping ends, and 75X coverage.

#### Oxford Nanopore long read

For nanopore sequencing, a protocol similar to previous conifer genome sequencing projects was used (Scott et al. 2020; Neale et al. 2022). Because sequencing by the Oxford Nanopore Technologies (ONT) platform requires more DNA per run and cannot be amplified to maintain read length, needle tissue was used for DNA extraction and sample preparation. High molecular weight DNA was extracted following the protocol described in Workman et al. (2018). Briefly, tissue was ground in liquid nitrogen with a mortar and pestle for 20 minutes to properly disrupt tissue. This is followed by lysis in a nuclear isolation buffer (NIB) containing spermine, spermidine, triton, and β-mercaptoethanol in a 50mL Falcon tube, rotating at 4C for 15 minutes. The resulting lysed sample is filtered through a steriflip, then centrifuged 1900 x g for 20 minutes at 4C. The supernatant was decanted, and the pellet resuspended in 1mL of NIB with a paintbrush. The resuspension was brought to a total volume of 15 mL NIB and centrifuged 1900 x g for 10 minutes at 4C. These steps were repeated (discard supernatant, resuspend pellet and wash) until the supernatant was clear, usually 2-3 times. The final pellet was resuspended into 1 mL 1X HB buffer per gram of initial tissue. Nuclei can then be spun at 7000xg for 5 minutes, supernatant removed and pellets snap frozen in liquid nitrogen and stored at −80C for later DNA extraction.

Extracted nuclei were then lysed and gDNA precipitated using the Circulomics Nanobind Plant Nuclei Big DNA kit, alpha version (EXT-PLH-001). DNA was sheared to 25kb with the Megaruptor 2, and library preparation was performed according to the ligation sequencing kit (LSK109, ONT). Then 1 μg of purified genomic DNA was input into the ligation sequencing kit (LSK108-LSK109, ONT). Samples were sequenced on R9.4 flowcells on either the minION or PromethION and then base-called using guppy 4.011-5.0.13 depending on time of sequencing.

### Assembly

The initial contig assembly utilized both ONT and Illumina data with a hybrid approach, where the ONT reads were first corrected using the Illumina reads, and then the corrected reads were assembled. Following the strategy used in our previous work assembling loblolly pine (Zimin et al. 2014) and other conifers, the whole-genome Illumina libraries were prepared from haploid megagametophyte tissue collected from a single seed. This resulted in reduction of the effective genome size, lowered the resource requirements on the hardware, and produces a more accurate assembly overall.

The contigs were assembled with MaSuRCA version 4.0.6 (Zimin et al. 2017). MaSuRCA uses the “super-reads” technique to compress Illumina reads by turning high-coverage Illumina paired end reads into low (2x to 3x) coverage of much longer super-reads that preserve the information contained in the Illumina reads. The super-reads were then used to error-correct the ONT reads, essentially producing mini-assemblies for each ONT read by using the ONT read as template. This process yields highly accurate “mega-reads,” with typically one or a few mega-reads corresponding and covering each nanopore read. The mega-reads are then assembled with a modified version of the Flye assembler (Kolmogorov et al. 2019).

Table 1 lists the data that were used for initial contig assembly of the whitebark pine genome along with the sizes of intermediate super-reads and mega-reads. The Flye assembler has an internal limitation of total input sequence of 549 Gbp. To stay within this limit, a subset of the longest mega-reads was used as input to the Flye assembler. The Flye assembly process was also modified. The assembler was interrupted after the initial contig (called disjointig in the Flye paper terminology) building stage to skip the initial contig consensus. This was necessary because the Flye consensus algorithm would otherwise attempt to create a >50 TB file of alignments of mega-reads to the contigs and eventually fail on data of this size. The consensus step was not needed because the mega-reads supplied to Flye were highly accurate. After skipping the consensus, the assembly continued with the repeat resolution and scaffolding steps. This process is automated in MaSuRCA (as of version 4.0.7 and higher). The new versions automatically perform the necessary steps when the detected genome size is over 10 Gbp. The statistics for this initial contig assembly (v0.1) are listed in Table 2.

**Table 1.** Quantitative statistics of the initial sequencing data and intermediate processed reads. Super-reads were produced from Illumina reads. Mega-reads were built from super-reads using ONT reads as templates. Each ONT read yielded one or several nonoverlapping mega-reads.

|  | Total sequence (bp) | Count | N50 size (bp) |
| --- | --- | --- | --- |
| *Original sequence data* |  |  |  |
| Illumina reads | 2,511,282,622,124 | 18,120,442,068 | 151 |
| Nanopore reads | 571,078,527,938 | 53,691,131 | 19,989 |
| Nanopore ultralong reads | 322,256,363,438 | 15,170,331 | 42,785 |
| *Derived data* |  |  |  |
| Super-reads | 85,674,725,052 | 163,046,437 | 1,228 |
| Mega-reads (subset used for Flye assembly) | 548,999,989,868 | 23,818,550 | 23,140 |

**Table 2.** Quantitative statistics of the intermediate and final assembly steps. The initial assembly (v0.1) was performed with the MaSuRCA assembler. That initial assembly was followed by scaffolding with SAMBA and polishing with POLCA to yield assembly v0.9. That assembly was filtered for redundancy, scaffolded by the HiRise scaffolder and then super-scaffolded into chromosome-sized scaffolds with the ALLMAPS software, followed by SAMBA gap closing and polishing with JASPER to yield the final assembly (v1.0). N50 contig size decreased going from v0.9 assembly to v1.0 assembly because HiRise scaffolder breaks contigs that are inconsistent with the HiC data. Consensus quality was evaluated with POLCA software.

| Assembly version | Total sequence (bp) | Number of contigs | N50 contig size (bp) | Number of scaffolds | N50 scaffold size (bp) | Consensus quality (%) |
| --- | --- | --- | --- | --- | --- | --- |
| v0.1 | 26,961,471,748 | 194,849 | 389,205 | 194,178 | 397,606 | 99.97 |
| v0.9 | 27,687,627,594 | 101,182 | 727,847 | 100,511 | 735,520 | >99.999 |
| v1.0 | 27,605,955,854 | 92,740 | 537,007 | 34,176 | 2,005,774,401 | >99.999 |

The initial contig assembly was followed by long-read contigging/scaffolding with SAMBA scaffolder (Zimin and Salzberg 2022). The original, uncorrected ONT reads that were 10 Kbp or longer were used for SAMBA scaffolding. Some of these reads may have been omitted in the contig assembly because of the input size limitation of the Flye assembler. The first iteration of SAMBA was very conservative, requiring ONT reads to match for a minimum of 9 Kbp to the ends of two contigs to join them. In the second iteration, that requirement was reduced to 4 Kbp. The scaffolder merges contigs and computes the consensus sequence filling the gap using the sequence of multiple ONT reads spanning the gap. Therefore the “patches” that filled the gaps may have a higher error rate.

The final step of the contig assembly was polishing the assembly with Illumina data in two passes using POLCA (Zimin and Salzberg 2020). The initial quality of the contigs after the SAMBA scaffolding was estimated to be 99.988% or QV39. After two rounds of POLCA polishing, the consensus quality was 99.999% or QV50, corresponding to an estimated error rate of 1 per 100,000 bases. These steps resulted in assembly v0.9, with statistics shown in Table 2.

Next, the contigs were scaffolded with OmniC reads (a variant of the HiC proximity ligation technique) with the HiRise scaffolder (Putnam et al. 2016) at Dovetail Genomics (now part of Cantata Bio). After the HiRise scaffolding, redundant duplicate contigs were identified. These exist because assemblers frequently leave extra copies of repeats or extra copies of alternative haplotype sequences already represented in the contigs as short contigs in the assembly. All 20,661 “short” contigs that were shorter than 10,000 bp were aligned to the rest of the assembly with nucmer aligner. These contigs contained 96,950,513 bp of sequence with N50 of 5,302 bp. Any contig that was shorter than 10,000 bp and that aligned to an interior of another contig with >95% identity over >95% if its length was removed from the set of short contigs. The remaining 1,371 short contigs containing 6,341,804 bp were added back to the assembly.

Following scaffolding with the OmniC data and redundancy filtering, two linkage maps (Weiss et al. 2020; De La Torre 2023) for the closely related sugar pine genome were utilized to super-scaffold the assembly to obtain chromosome-sized scaffolds. The two maps had a total of 7,767 markers (mostly short sequences). Of these markers, 2,959 mapped uniquely to the whitebark pine scaffolds. ALLMAPS software (Tang et al. 2015) was utilized to produce chromosome-sized scaffolds using the alignments of markers to the scaffolds and marker positions in the map. The two final steps following the scaffolding were additional gap closing with the SAMBA tool (Zimin and Salzberg 2022) using the ONT reads followed by polishing with the JASPER polisher (Guo et al. 2023) that used the Illumina data. This additional polishing was needed because in the places where gaps in the scaffolds were filled, consensus computed only from the ONT reads that spanned these gaps would have resulted in low-quality sequence. The statistics of this final assembly (v1.0) are listed in Table 2.

### Annotation and comparative genomics

#### Transcriptomic evidence

A combination of public RNA-Seq (Illumina PE) data from mixed tissue types was employed for the first stage of annotation (PRJNA703422, PRJNA352055). Illumina short reads were aligned to the v0.9 reference genome with HISAT2 v2.2.1, including the following flag to accommodate long-introns --max-intronlen 2500000 (Kim et al. 2019). All libraries with mapping rates that exceeded 95% alignment and contained a minimum of 20 million reads were retained for evaluation (Table S1). In addition, a set of recently de novo-assembled libraries (Illumina NovaSeq 150 bp PE), from needle tissue of six individuals from a single half-sib family from Shadow Lake 39, Mount Rainier National Park collected and flash frozen by the USDA Forest Service Dorena Genetic Resource Center (BioProject PRJNA933606) were used as further evidence (Table S2). These were assembled with the Oyster River Protocol (ORP) workflow (v2.2.5; MacManes 2018), a combined pipeline that works with Trinity v2.9.1 (Haas et al. 2013), rnaSPAdes v 3.13 (Bushmanova et al. 2019), and TransABySS v2.0.1 (Robertson et al. 2010) assemblers to generate a single reference assembly. This assembly was subsequently clustered at 90% with USearch (v9.0.2132; Edgar 2010), frame-selected with Transdecoder (v.5.5.0; https://github.com/TransDecoder/TransDecoder), and filtered with eggNOG (v4.1; Huerta-Cepas et al. 2019). This transcriptome was further filtered for short fragments (<300bp) with SeqKit (v2.2.0; Shen et al. 2016) and aligned to the v0.9 genome reference via Minimap2 ([-ax splice:hq -uf]; v2.24; Li 2018). Secondary alignments produced by Minimap2 were removed via SAMtools (v1.9; Danecek et al. 2021).

#### Structural annotation of the v0.9 genome

Initial assessment of the v0.9 reference genome was conducted with BUSCO v5.2.2 with the embryophyta database (odb10; Manni et al. 2021). Subsequently, repeat sequences were identified de novo with a combination of self-to-self comparisons and structural identification with RepeatModeler v2.01 (Flynn et al. 2020). The twice soft-masked genome was used as input to Braker v2.1.5 as well as the aligned RNA-Seq reads from NCBI (Brůna et al. 2021). The set of predicted proteins was filtered with eggNOG v5.0.2 and evaluated with QUAST v5.2.0 (Gurevich et al. 2013) and BUSCO (embryophyta). In parallel, StringTie2 v2.2.1 was run using different sets of transcriptomic input (Kovaka et al. 2019). The first gene space assembly utilized only the HISAT2 aligned short reads as input to StringTie2, while the second assembly was run in hybrid mode including both the HISAT2 alignments and full-length assembled transcripts assembly aligned to the genome with Minimap2 (Li 2018). Protein-coding sequences were generated from all StringTie2 runs with Gffread v0.12.1 (Pertea and Pertea 2020) and frame-selected and filtered with Transdecoder v5.5.0 and eggNOG v5.0.2. Supplemental Transdecoder scripts were used to obtain the coordinates of the frame-selected transcripts in the context of the genome. Transcripts that did not have a corresponding genome alignment after this filtering step were removed from the final coding sequence and protein sequence files. The final proteins were evaluated with BUSCO (embryophyta), EnTAP v0.10.8, and AGAT v1.0.0 (Hart et al. 2020; Dainat 2022). EnTAP was run as a reciprocal BLAST search to estimate alignment rate at 50/50 coverage between the query sequence and target databases (NCBI’s RefSeq v208 and UniProt). AGAT was employed to provide basic filtering for structural anomalies and quantify statistics regarding structural aspects of the protein-coding regions (Dainat 2022). After EnTAP annotation, transcripts without a similarity search or eggNOG match were scanned for protein domains using InterProScan, and those lacking any identifiable protein domains were removed.

#### Annotating the v1.0 genome

Initial assessment of the v1.0 genome was conducted with BUSCO v5.4.5 with the embryophyta database. Following the generation of v1.0 of the genome assembly, the annotation lift-over tool Liftoff was used to map the v0.9 annotation onto the v1.0 genome (Shumate and Salzberg 2021). The parameter -*a 0.8* was used to specify that the minimum coverage of the mapped genes should be at least 80%. The parameter *-s 0.8* was used to specify that the minimum sequence identity of the mapped genes should be 80%. After transferring the annotation, Gffread was used to extract the coding sequences and proteins from the genome. The final proteins were evaluated with BUSCO (embryophyta), EnTAP, and AGAT.

#### NLR identification on the v0.9 genome

Three methods were utilized to generate a more complete representation of potential nucleotide-binding and leucine-rich repeats (NLRs) in whitebark pine: InterProScan, RGAugury, and NLR-Annotator. NLRs were identified from a de novo-assembled transcriptome, whole genome scanning, and from the genome annotation to provide comparison across the available genomic resources.

InterProScan v5.35-74.0 and RGAugury v1.0 identified NLRs from the protein sequences of the genome annotation through protein domain scanning (Li et al. 2016; Paysan-Lafosse et al. 2023). Interproscan was used to identify the NB-ARC, TIR, CC, RPWB, and LRR domains using the Pfam, Gene3D, SUPERFAMILY, PRINTS, SMART, and CDD databases. The GFF3 file produced by InterProScan was filtered using a custom python script to remove all entries without at least one NLR domain, to speed up the identification and classification steps downstream. Custom R scripts were employed to identify the NLRs and classify them into their subfamilies based on the N-terminal domain. Those with a TIR domain are TNLs, those with a coiled-coil (CC) domain are CNLs, and those with an RPW8 domain are RNLs. Subfamilies included both complete NLRs (containing N-terminal, NBARC, and LRR domains) and those missing just the LRR domain. Sequences without an N-terminal domain (NBARC only and NBARC-LRR) were considered unclassified. The RGAugury pipeline is quite similar, but it first implements a filtering step based on sequence similarity to the Resistance Gene Analog database before performing domain scanning with InterProScan. RGAugury was better able to identify CNL type NLRs than InterProScan which struggled to identify the CC N-terminal domain.

NLR-Annotator v2.0 was used to identify potential NLRs directly from the genome sequence using NLR-associated DNA motifs (Steuernagel et al. 2020). From the genome annotation, genes overlapping at least 80% of the predicted NLRs based on the NLR-Annotator boundaries were selected as potential NLRs with BEDTools v2.29 (Quinlan and Hall 2010). Custom R scripts were employed to combine NLR annotation results from the three methods and identify which annotations were unique to each method. To reintroduce gene models from the BRAKER annotation, gene predictions that overlapped at least 90% of the boundaries of a complete NLR (CNLs and TNLs) were retained and included in the primary genome annotation.

## Results and Discussion

### Sequencing

Previously developed sequencing methods to analyze other conifers (Scott et al. 2020) were used to generate a combination of short (Illumina) and long (ONT) sequencing data in whitebark pine. This fusion of technologies brings together the advantages of both approaches: leveraging long nanopore reads to span repetitive sequences commonly found in conifers producing a highly contiguous genome assembly. Although the error rate of nanopore sequencing is steadily improving, it still poses challenges for the final assembly. By integrating these long reads with highly accurate, albeit shorter, Illumina reads, a more precise assembly was produced while maintaining a high level of contiguity.

First, short-read Illumina sequencing data were generated from DNA of a megagametoyphyte. The haploid megagametophyte DNA precludes the typical difficulties associated with diploid DNA and natural genetic variation between alleles. From this DNA, ∼2.5Tb of sequence was generated for an estimated ∼100X coverage (Table 1).

It has previously been found that short-read sequencing, especially in conifers, results in low contiguity as the highly repetitive areas typical to these genomes are impossible to assemble with short reads alone. Complicating this issue, existing long-read sequencing methods require relatively large amounts of DNA, and achieving the long read length precludes the use of PCR. As an alternative, high-molecular weight (HMW) genomic DNA from needle tissue was extracted from the same tree (Workman et al. 2018). Using a combination of cryogenic tissue grinding and nuclei extraction, high quality DNA was obtained, which was then subjected to either long-read (N50 20kb, 571Gb, ∼23X) or ultralong-read (N50 42.8kb, 322Gb, 13X) nanopore sequencing.

### Assembly

The MaSuRCA assembler transformed the Illumina reads into super-reads by first constructing a k-mer graph from k-mers (k=99 here) found in the Illumina reads. The k-mers are the nodes in the k-mer graph, and exact overlaps of k-1 bases between k-mers are the edges. The super-reads technique then uses the graph to extend each Illumina read in 5’ and 3’ directions as far as possible, as long as the extension is unambiguous, i.e., there are no branches in the k-mer graph. The extended read is called a super-read. Many Illumina reads extend to the same super-read. The technique transforms the Illumina reads into super-reads that contain the same information, however there are many fewer super-reads and they are much longer. Table 1 shows that the super-read transformation turned over 18 billion 151 bp Illumina reads into about 163 million super-reads. Half of the sequence in the super-reads was in sequences of 1228 bp or longer. MaSuRCA then used super-reads to correct the ONT reads by building mini-assemblies of overlapping super-reads for each ONT read. These mini-assemblies are produced using the ONT reads as templates, and they are called mega-reads. Mega-reads are long and they have a very low error rate, less than 0.5%. The mega-reads algorithm resulted in producing about 24 million mega-reads with N50 size of 23,140 bp. The MaSuRCA assembly (v0.1) (Table 2) was followed with scaffolding with SAMBA and polishing with POLCA, resulting in assembly v0.9 (Table 2). The v0.9 assembly was then scaffolded with HiRise with OmniC data and super-scaffolded with ALLMAPS using the alignments of markers to the scaffolds and marker positions from the sugar pine map. Fig. 1 shows the alignment of the markers from the sugar pine maps to the whitebark pine super-scaffolds. Some discrepancies between the scaffolds and the map were observed in chromosomes 1, 2, 3, 5, 6, and 11. These discrepancies could be due to interchromosomal rearrangements between the sugar pine and whitebark pine genomes. However, they could also be due to mis-assemblies in the scaffolds of whitebark pine, which cannot be resolved with the currently available data. Scaffolding with ALLMAPS resulted in 24,069,114,767 bp of sequence anchored to the chromosomes of which 23,671,235,725 bp was also oriented. Additional gap-closing was then applied to the scaffolds with the SAMBA tool that used original uncorrected ONT reads to fill gaps in the scaffolds. SAMBA closed 1,484 gaps in the assembly, adding an 9,065,412 bp of sequence to the assembly. Finally, the JASPER tool was applied to polish the assembly with the Illumina reads. The final polished assembly (v1.0) (Table 2) has an error rate of less than 1 error in 100,000 bases and it contains 27,605,955,854 bp of sequences in 34,176 scaffolds with N50 contig size of 537,007 bp. Approximately 87.2% (24,072,309,274 bp) of total sequence was placed on the twelve chromosomes.

**Fig. 1.**
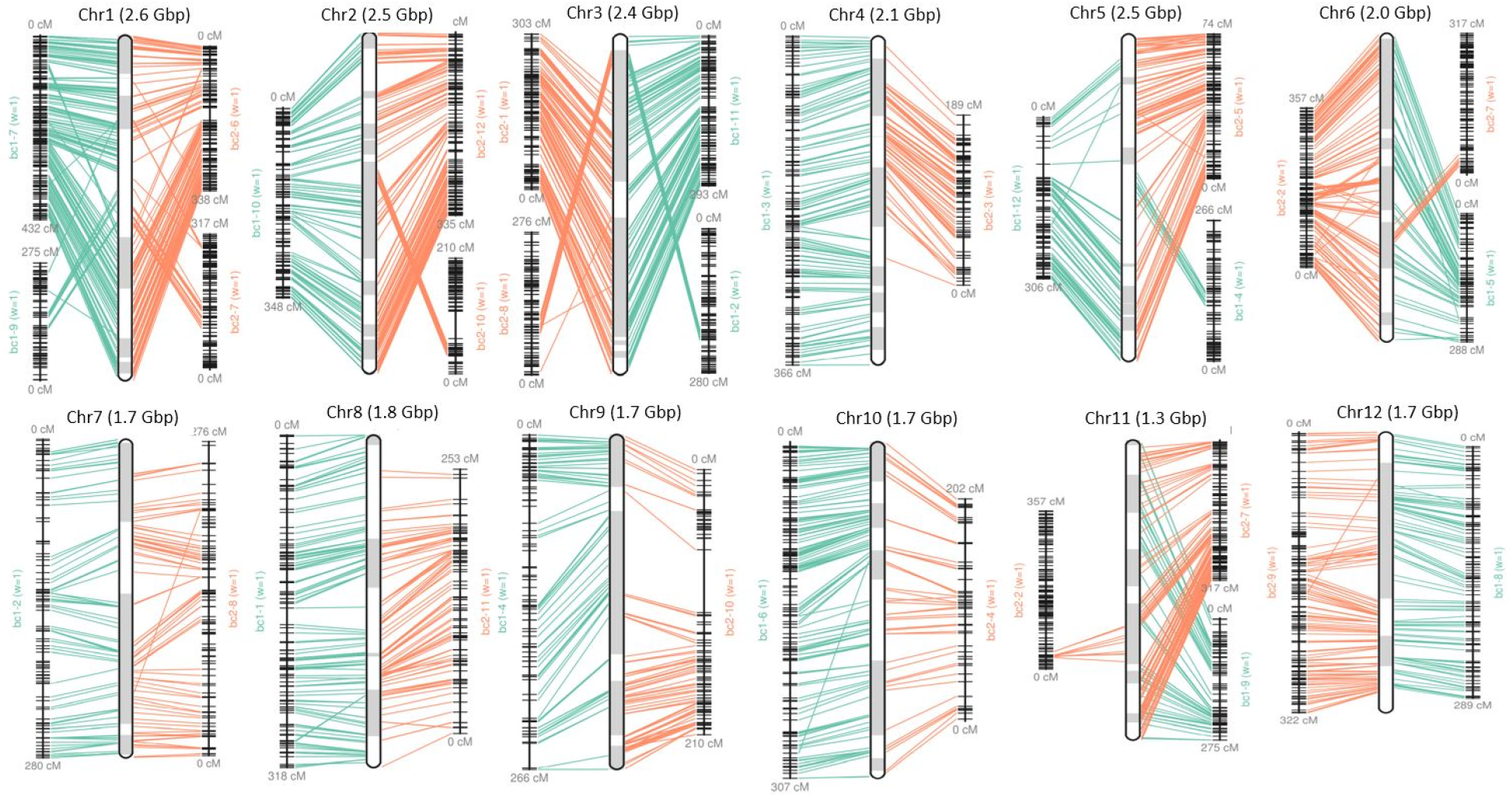
Alignment of the sugar pine linkage map markers to the whitebark pine super-scaffolds. The individual chromosome plots are produced by the ALLMAPS software. The vertical bars in the middle of each panel represent the chromosomes. The two maps are shown on the two sides of each chromosome with marker alignments shown in green or orange. The individual scaffolds within the chromosome are shown in in white or gray shading inside the middle vertical bar.

### Annotation

#### Identifying and masking repetitive regions

Prior to the alignment of the transcriptomic short reads, repeat identification with RepeatModeler generated a custom library of 2576 unique repeat sequences, of which 558 could be classified (long terminal repeat (LTR): 502 (Copia: 152, Gypsy: 347); LINE: 25; Others: 33) (Table S3). This repeat library was used with RepeatMasker to softmask 77.6% of the genome sequence (Table 3). The overall repetitive content was comparable to *Pinus taeda* at 74% and *P. lambertiana* at 79% (Stevens et al. 2016). The majority of the repetitive elements were LTRs, which comprised almost 42% of the genome, and roughly 32% of the genome was unclassified repetitive sequences (Table S3). The high proportion of unclassified elements is likely due to RepeatModeler being unable to classify many of the repeats in the generated custom repeat library that was used to mask the genome, as LTRs comprised only 55% of the repeat content in whitebark pine where they usually contribute around 70% of the TE content in conifers (De La Torre et al. 2014; Stevens et al. 2016; Fujino et al. 2023).

**Table 3.** Statistics on the structural annotation of the whitebark pine reference genome assembly.

|  | **v0.9 assembly** | **v1.0 assembly** |
| --- | --- | --- |
| *Completeness* |  |  |
| Genome BUSCO v5 | C:55.3% [S:45.5%, D:9.8%], F:24.3%, M:20.4% | C:65.5% [S:57.1%, D:8.4%], F:19.1%, M:15.4%, n:1614 |
| Annotation BUSCO v5 | C:70.6% [S:43.2%, D:27.4%], F:15.1%, M:14.3% | C:67.0% [S:40.7%, D:26.3%], F:14.4%, M:18.6%, n:1614 |
| *Protein-coding genes* |  |  |
| Total number of genes | 27,010 | 25,362 |
| Number of single-exon genes | 4,836 | 4,689 |
| Number of multi-exon genes | 22,174 | 20,673 |
| Mono:multi ratio | 0.22 | 0.22 |
| Total number of transcripts | 47,911 | 45,095 |
| Transcript N50 | 1578 bp | 2571 bp |
| Longest intron | 1.39 Mbp | 1.39 Mbp |
| Average number of exons | 6 | 6 |
| *Functional annotation* |  |  |
| EnTAP annotation rate (gene family) | 92.8% | 83.65% |
| EnTAP annotation rate (sequence similarity search) | 86.5% | 77.50% |
| *Repeat detection* |  |  |
| Softmasked % (LTR %) | 77.6% (55%) |  |

#### RNA sequence data for annotation and transcriptome assembly

A total of 12 Illumina RNA-Seq libraries (Table S1) were mapped to the whitebark pine reference v0.9 genome following quality control. The final set of selected Illumina libraries ranged from 23.9 to 66.9 M reads and aligned well to the reference (94.8 to 96%). These alignments were used with BRAKER and StringTie2 to generate the draft genome assembly (Kovaka et al. 2019; Brůna et al. 2021). Two RNA-seq libraries (SRR13823648 and SRR13823649) generated from megagametophyte tissue were used for a de novo transcriptome assembly that was utilized for NLR annotation (Table S1). This transcriptome assembly consisted of 37,586 transcripts and had a BUSCO (embryophyta) completeness of 92.6% [S:88.8%, D:3.8%].

#### Preliminary annotation of the v0.9 genome

The preliminary protein-coding predictions generated from BRAKER amounted to an overestimate with 636.6K initial models (BUSCO: C:45.0% [S:35.6%, D:9.4%]; Table S4). Following basic gene family level filtering with eggNOG, a total of 219.5K transcripts (BUSCO: C:45.0% [S:35.7%, D:9.3%]; N50:1164; longest Intron:140 Kb) were retained. The short reads processed by StringTie2 produced a total of 63,123 transcripts (BUSCO: C:70.9% [S:43.3%, D:27.6%]; N50 2281 bp). These transcripts were filtered via Transdecoder/eggNOG, leaving a total of 48,567 transcripts (BUSCO: C:70.5% [S:43.2%, D:27.3%]; N50 1578 bp; Longest Intron: 1.39 Mb).

To improve upon challenges associated with short-read alignment against the complex and repetitive conifer genome, the de novo-assembled transcripts resulting from an independent transcriptomic sampling were aligned at 71% to the genome with Minimap2 (Table S2). These transcripts were then used as long-read input for a hybrid long- and short-read transcriptome assembly using StringTie2. The hybrid run of StringTie2 generated a total of 62,936 transcripts (BUSCO: C:71.4% [S:37.4%, D:34.0%]; N50 1807). These gene models were filtered via Transdecoder/eggNOG, resulting in a total of 45,380 transcripts (BUSCO: C:70.4% [S:45.7%, D:24.7%]; N50 1515; Longest Intron: 1.02 Mb; Table S4).

As an additional metric for completeness, the de novo-assembled transcripts were aligned to the reference genome independently, resulting in a total of 66,233 unique alignments (BUSCO: C: 88.50% [S:46.60%, D:41.90%]; N50 2217; Table S4). These alignments represent variation and gaps and do not directly translate to viable protein-coding models but can provide a benchmark for completeness.

#### Filtering the v0.9 genome annotation

The Transdecoder/eggNOG-filtered StringTie2 short-read predictions were selected as the best overall annotation. This annotation was further refined by removing transcripts without an EnTAP similarity search or eggNOG annotation that also lacked any protein domains identified using InterProScan, reducing the annotation by 683 genes. An additional 27 complete NLR genes identified from the genome using NLR-Annotator, and overlapping a gene model generated by BRAKER, were added to the annotation (Table S5). This final set consisted of 27,010 genes and represented a total of 47,911 transcripts. The annotation had a BUSCO completeness of 70.6% [S:43.2%, D:27.4%], and an EnTAP similarity search annotation rate of 85%, and the longest intron recorded was 1.39 Mb in length (Table 3). The annotated gene space of whitebark pine is larger and more representative than those of *Pinus lambertiana* and *P. taeda*, which contained 13,936 and 9,024 high confidence genes with BUSCO completeness of 53% and 30%, respectively (Stevens et al. 2016). The genome annotations of the spruce (*Picea*) species range from 35% to 49% completeness (Gagalova et al. 2022). More recent conifer genome assemblies report higher BUSCO completeness, such as *Sequoia sempervirens* at 65.5% completeness (Neale et al. 2022), *Pinus tabuliformis* at 84% (Niu et al. 2022), and *Cryptomeria japonica* at 91.4% (Fujino et al. 2023).

Compared to the 70.6% BUSCO completeness of the genome annotation, at the genome level, the whitebark pine genome accounted for only 55.3% BUSCO completeness using the embryophyta lineage, likely due to challenges associated with the predictions across long introns as well as the abundant pseudogenes and high repeat content (Table 3). This result is typical of conifer genomes and, despite this, the whitebark pine genome BUSCO completeness was slightly higher compared to that of several other recently assembled conifer genomes (sugar pine, spruce, coast redwood, and Chinese pine; Stevens et al. 2016; Gagalova et al. 2022; Neale et al. 2022; Niu et al. 2022, respectively).

#### Transferring the v0.9 genome annotation to the v1.0 genome

A total of 25,362 genes and 45,095 transcripts (from the original 27,010 genes) were transferred to v1.0. The BUSCO completeness of the v1.0 genome annotation was 67.0% [S:40.7%,D:26.3%] and the EnTAP similarity search annotation rate was 77%. This is a slight decrease from the 70.6% completeness and 85% annotation rate of the v0.9 genome annotation. The monoexonic to multiexonic gene ratio and longest intron length, 0.22 and 1.39 Mbp respectively, were equivalent.

#### NLR identification on the v0.9 genome

NLRs are a major class of disease resistance genes that recognize specific virulence factors. They have a characteristic domain structure with one of three canonical N-terminal domains, a nucleotide-binding domain, and a leucine-rich repeat domain. NLRs can be divided into subfamilies based on their N-terminal domain; TNLs contain a TIR domain, CNLs contain a coiled-coil domain, and RNLs contain an RPW8 domain (Van Ghelder et al. 2019). The combination of the three software methods used to identify NLRs from version 0.9 of the genome, before scaffolding, (InterProScan, RGAugury, and NLR-Annotator) was necessary to fully describe all three types of NLRs, as can be seen from the overlap between complete NLRs identified from the genome annotation by each method, or lack thereof (Fig. 2B). The three methods were able to independently identify the majority of TNL-type NLRs. Thirty-four TNLs were identified by all three methods, 11 were identified by domain scanning methods only, 11 were unique to NLR-Annotator, and one was unique to InterProScan. Here, the domain scanning methods performed equally well and the results of NLR-Annotator were a useful addition to the set of complete NLRs. InterProScan was necessary to identify the RNLs as the other two programs cannot identify the RPW8 domain and would otherwise identify these as NLRs missing an N terminal domain. RGAugury was necessary as NLR-Annotator and InterProScan were not as effective in identifying CNLs, and 7 complete CNLs were identified using RGAugury only.

**Fig. 2.**
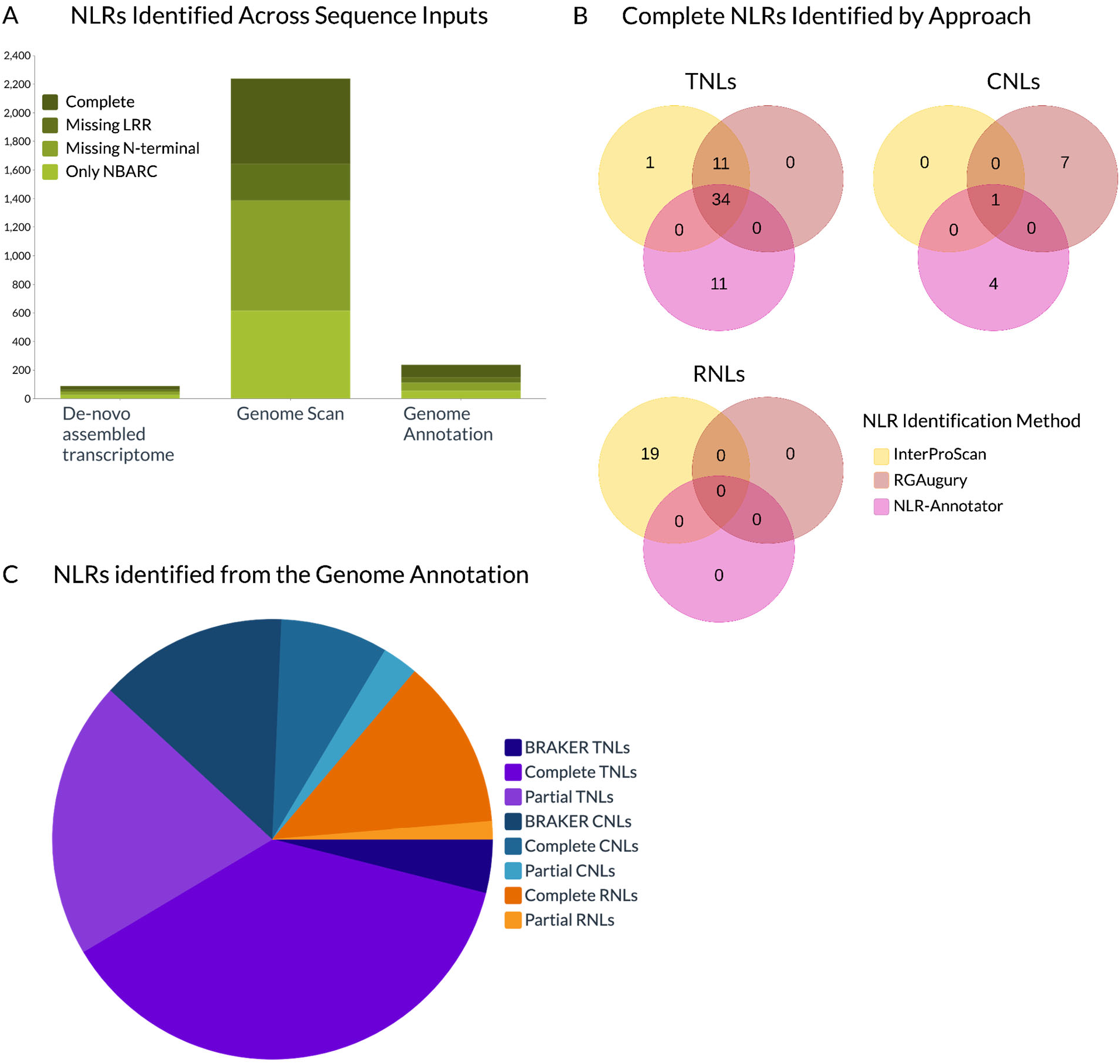
Results of NLR annotation methods. **A**) NLRs identified by input type; a de novo-assembled transcriptome, the genome sequence, and the genome annotation. The darkest green distinguishes complete NLRs, with progressively lighter shades to represent NLRs missing an LRR domain, NLRs missing an N-terminal domain, and NLRs identified only by the NBARC domain. **B**) Within the genome annotation, complete NLRs identified by each method and annotations with support from multiple methods. Yellow represents NLRs identified only using InterProScan. Coral/red represents NLRs identified using only RGAugury. Pink represents NLRs identified using only NLR-Annotator and supported by the genome annotation. **C**) Breakdown of total classified NLRs in the genome annotation with the addition of genes recovered from BRAKER and their contribution to the classes. Purple represents the TNL class, with the darkest shade representing complete TNLs that had support from BRAKER gene predictions, medium representing complete TNLs from the genome annotation, and lightest representing incomplete TNLs. Blue represents the CNL class, with the darkest shade representing complete CNLs that had support from BRAKER gene predictions, medium representing complete CNLs from the genome annotation, and lightest representing incomplete CNLs. Orange represents the RNL class which did not have any additional NLRs from BRAKER, with the darker shade distinguishing complete RNLs from the light incomplete RNLs.

Integration of the gene models from BRAKER that overlapped with complete NLRs from genome scanning was the greatest contributor to the CNL subfamily, contributing 21 complete CNLs out of a total of 33. The ratio of TNLs, CNLs, and RNLs are as expected in conifers, with TNLs being the largest class followed by CNLs, and RNLs being the smallest class (Van Ghelder et al. 2019). RNLs are also more abundant in conifers and some members of the Rosaceae compared to most land plants, which typically have ten or fewer RNLs (Van Ghelder et al. 2019). Without recovering complete CNL gene models from the ab initio BRAKER gene predictions of whitebark pine genes, there would have been double the number of complete RNLs identified compared to the number of CNLs. Based on NLR identifications in other conifers, the number of CNLs is generally two to three times as many the number of RNLs (Van Ghelder et al. 2019). In the giant sequoia (*Sequoiadendron giganteum*) genome, there were 53 complete CNLs and 17 complete RNLs identified (Scott et al. 2020), following the expected ratio. As the gene models in the whitebark pine genome annotation were dependent on RNA evidence, one contributor could be that CNLs were expressed at a lower level at time of sampling and therefore were not well represented in this genome annotation. The BRAKER pipeline utilizes ab initio gene prediction, which was able to predict the CNL gene models without direct RNA evidence.

NLRs were also identified from a de novo transcriptome assembly as well as by directly scanning the v0.9 genome as a comparison to the v0.9 genome annotation. A total of 89 potential NLRs were identified from the de novo-assembled transcripts using InterProScan, RGAugury, and a modification of the motif-finding portion of NLR-Annotator (Fig. 2A; Table S5). Of the 89 NLRs, 24 were considered complete, meaning they contained an N-terminal domain, an NB-ARC domain, and an LRR domain. From the genome annotation, 238 potential NLRs were identified using InterProScan, RGAugury, and the results of the NLR-annotator scan on the whole v0.9 genome. Gene annotations with the gene model overlapping with at least 80% of the predicted NLR boundaries from the genome scan were identified as potential NLRs. These results were combined with those from the domain scanning methods, resulting in a total of 88 complete NLRs. About three times as many NLRs could be identified using the v0.9 genome annotation compared to the transcriptome with a noticeable increase in completeness (Fig. 2A; Table S5). This likely results from a combination of challenges associated with de novo transcriptome assembly, such as fragmentation and fewer total number of transcripts (Transcriptome: 37.5 K transcripts, N50: 744 bp; v0.9 Genome annotation: 47.9 K transcripts, N50: 1578 bp). The genome annotations reflect a combination of more transcriptomic input as well as transcripts assembled using the genome as guidance, which likely provides a more accurate representation of the gene space.

NLRs have been extensively studied in angiosperms, allowing for the creation of RefPlantNLR which contains 481 NLRs with representatives from species across 31 genera, many of which have been experimentally validated (Kourelis et al. 2021). In comparison, NLRs have been cataloged in eight conifer species across six genera using primarily transcriptomic resources (Van Ghelder et al. 2019; Scott et al. 2020; Bondar et al. 2022; Ence et al. 2022). A better understanding of NLRs in conifer species would help to explore the mechanisms of disease defense in conifers and provide candidates for disease resistance genes. The InterProScan method of NLR identification was adapted from a prior study that identified between 338 and 725 NLRs across seven conifer species transcriptome assemblies, including two species of *Pinus* (Van Ghelder et al. 2019). At the lower end, this is three times the number of NLRs found in the whitebark pine transcriptome and more than the NLRs found in the genome annotation, indicating that the input may have an impact on the NLRs that are able to be identified. Utilizing a combination of NLR identification methods did improve the ability to identify NLRs compared to using InterProScan alone, and they were especially important for identifying the CNL class of NLRs.

Using NLR-Annotator to scan the v0.9 genome directly, 2,239 potential NLRs were identified, of which 595 were complete (Fig. 2A; Table S5). However, only 151 of the 2,239 NLRs overlapped with a gene from the v0.9 genome annotation and 54 of them were considered complete. The partial NLRs identified through genome scanning without an overlapping genome annotation are most likely pseudogenes and nonfunctional NLRs. NLRs are under rapid evolution and often undergo tandem duplications and rearrangements or recombinations, and pseudogenes with significant deletions or missing domains can accumulate (Marone et al. 2013). Partial NLRs identified by NLR-Annotator that have support from the genome annotation could be from incomplete transcripts or truncated gene models, as the annotation only included gene models with RNA evidence before genes were recovered from BRAKER. Twenty-one complete CNLs and six complete TNLs identified by NLR-Annotator that were not in the genome annotation but were supported by gene models from BRAKER generated from the v0.9 genome were included in the overall v0.9 genome annotation, resulting in a total of 265 candidate NLRs, of which 116 were complete (Fig. 2C; Table S5). This is far less than the 595 complete NLRs predicted using NLR-Annotator to scan the genome directly. Some “complete” NLRs may have only recently become nonfunctional and therefore less fragmented. Others may be real NLRs that were not expressed in any of the RNA-seq samples used for the annotation or predicted via ab initio methods. For comparison, 375 NLRs were identified in the giant sequoia reference genome examining the intersection between NLR-Annotator genome-scan predictions and the genome annotation. These gene models were primarily composed of BRAKER predictions that were supplemented with full-length transcript alignments. The pseudo-chromosomal assembly of giant sequoia also made it possible to identify the uneven distribution of the NLRs throughout the genome (Scott et al. 2020). With improved contiguity and completeness of both the genome and annotation, more NLRs are likely to be identified in whitebark pine.

This first in-depth classification of these elements in whitebark pine provides candidates for genes contributing to quantitative disease resistance against white pine blister rust. NLRs have been identified as candidate genes for major disease resistance loci in *P. lambertiana* (*Cr1*) (Stevens et al. 2016)*, P. monticola* (*Cr2*) (Liu et al. 2013), *P. strobiformis* (*Cr3*) (Liu et al. 2022b), and *P. flexilis* (*Cr4*) (Liu et al. 2021). Some NLRs have been identified as candidates for quantitative disease resistance in *P. lambertiana* (Weiss et al. 2020). A more recent study of the *Cr1* locus developed the marker Cr1AM1 to identify SNPs in the region associated with *Cr1* within the *P. lambertiana* genome (Wright et al. 2022). In version 1.5 of the *P. lambertiana* genome, variants of this marker aligned with greatest identity to two locations within the 6.3 Mb fragscaff scaffold_6044. These alignments did not directly overlap with any annotated genes on the scaffold. Though the alignment suggests that *Cr1* is an intergenic locus, it may be affecting the activation or expression of nearby genes resulting in the disease resistance response. There were 18 genes located on this scaffold and seven were NLRs genes, providing candidate genes for disease resistance to white pine blister rust in *P. lambertiana*.

Among the North American white pines, *P. albicaulis* has demonstrated the greatest variation in resistance to white pine blister rust (WPBR) across its extensive range. To date, patterns of major gene resistance have not been identified, suggesting a different mechanism from that of *P. lambertiana*, *P.strobiformis*, and *P.flexilis* (Liu et al. 2022a). Despite this, the putative Cr1AM1 marker sequence was aligned to the *P. albicaulis* v0.9 genome assembly. The best alignment, recorded at 93%, was to scaffold_64902 of length 327 Kb with no annotated genes. Five genes were identified on scaffold_64902 from the BRAKER annotation, but none of these putative genes were homologous to or aligned near the genes annotated on scaffold_6044 in *P. lambertiana*. The scaffold identified in the *P. albicaulis* genome also exhibits little sequence similarity to the scaffold identified in *P. lambertiana*. Further studies are needed to provide a comprehensive representation of the NLR space in *P. albicaulis* and identify specific candidates for improved disease resistance.

## Summary and Conclusion

This paper reports the first important step in developing genomic technologies that can be employed to more efficiently and rapidly identify genetic resources that can be used in the restoration of the threatened whitebark pine: a well-assembled and annotated reference genome sequence. As this core research team has done previously (generating reference genome sequences for five other conifer species [*Pinus taeda*, Neale et al. 2014; *P. lambertiana*, Stevens et al. 2016; *Pseudotsuga menziesii*, Neale et al. 2017; *Sequoiadendron giganteum*, Scott et al. 2020; and *Sequoia sempervirens*, Neale et al. 2022]), DNA from a single tree was used to generate the reference genome sequence. Comparison of genome size, gene number, and genome annotations among these genomes with that from whitebark pine reflects very strong similarity in gene and repetitive DNA content. However, at the phenotypic level, these conifers are quite different from each other in many ways (anatomy, morphology, life history, reproductive traits, adaptative traits, disease and insect susceptibility/resistance, etc.). These large phenotypic differences must be due in a large part to allelic variation among a common set of genes and the expression of these genes. Thus, research must now begin in whitebark pine to discover population-level allelic variation and variation in gene expression. As our team has done for all other conifer genome projects, we will now embark on the genome-wide association studies and environmental association studies to discover natural variation and investigate its relationship with the vast amount of phenotypic and adaptive variation in populations of whitebark pine. Discovery from studies of this nature will lead to development of applied genomic screening tools to be used in restoration programs.

## Data availability

The whitebark pine assemblies v0.9 and v1.0 were deposited at NCBI under BioProject PRJNA1034085. Annotation files for both v0.9 and v1.0 assemblies are available at the TreeGenes repository (https://treegenesdb.org/FTP/temp/P_albicaulis/).

## Supplementary material

**Table S1.** Summary statistics for transcriptomic evidence used for gene prediction (reads only)

| **SRA**<br>**accession** | **Bioproject** | **Tissue type** | **Raw reads**<br>**(M)** | **Trimmed reads**<br>**(M)** | **Mapping**<br>**rate** |
| --- | --- | --- | --- | --- | --- |
| SRR13823648 | PRJNA703422 | megagametophyte | 68.861096 | 66.916378 | 95.2 |
| SRR5368574 | PRJNA352055 | needle | 38.02665 | 37.16283 | 96 |
| SRR5368575 | PRJNA352055 | needle | 41.248962 | 40.279086 | 95.9 |
| SRR5368576 | PRJNA352055 | needle | 37.951822 | 37.951822 | 95.3 |
| SRR5368577 | PRJNA352055 | needle | 35.176356 | 34.236232 | 95.3 |
| SRR5368578 | PRJNA352055 | needle | 34.290692 | 33.466354 | 95.5 |
| SRR5368579 | PRJNA352055 | needle | 32.297144 | 31.417736 | 96 |
| SRR5368580 | PRJNA352055 | needle | 25.204748 | 24.604476 | 96 |
| SRR5368581 | PRJNA352055 | needle | 30.622742 | 29.958086 | 96 |
| SRR5368582 | PRJNA352055 | needle | 24.44717 | 23.868976 | 94.8 |
| SRR4786281 | PRJNA352055 | needle | 36.080828 | 35.09486 | 95 |
| SRR4786284 | PRJNA352055 | needle | 46.218548 | 44.478282 | 94.8 |

**Table S2.**
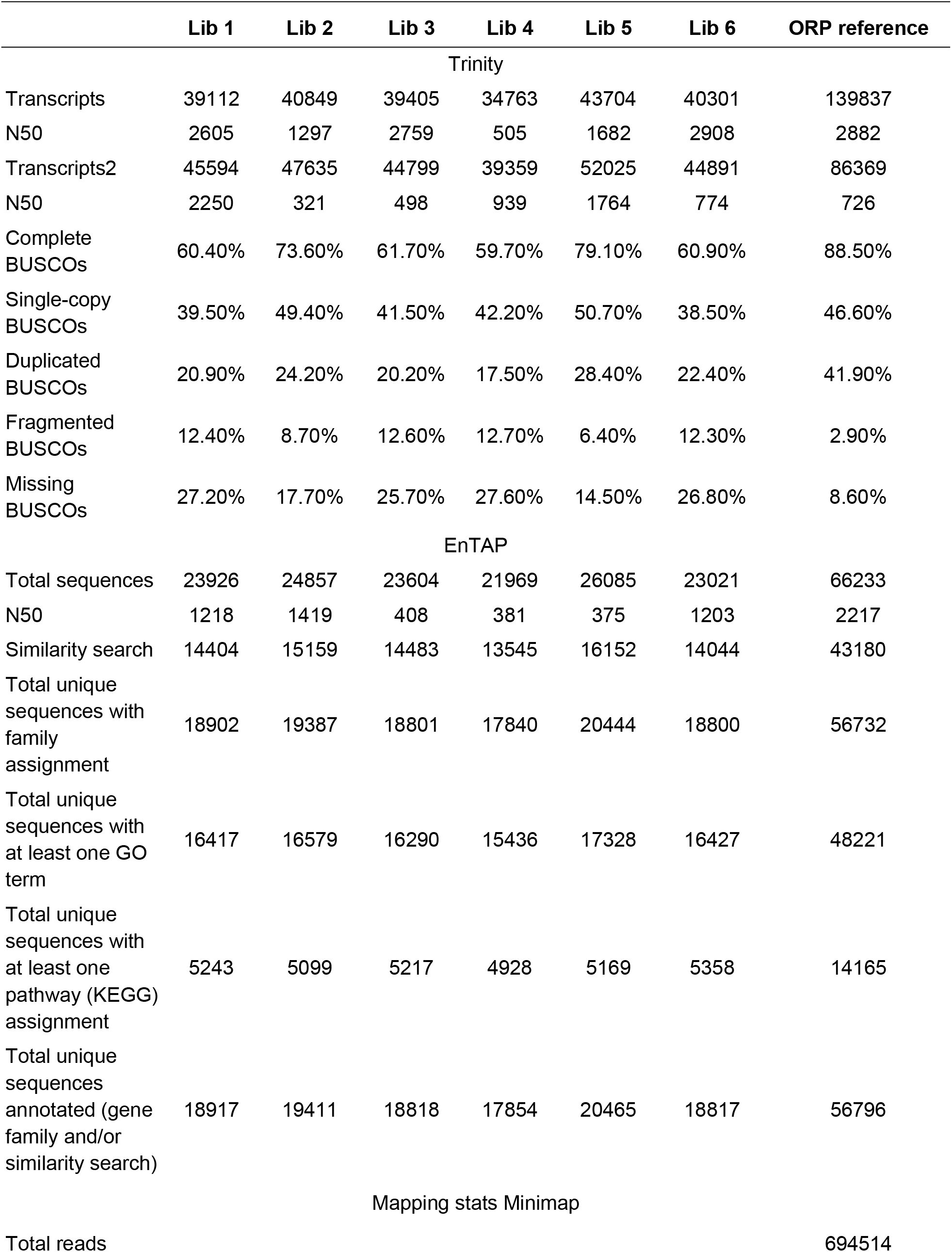
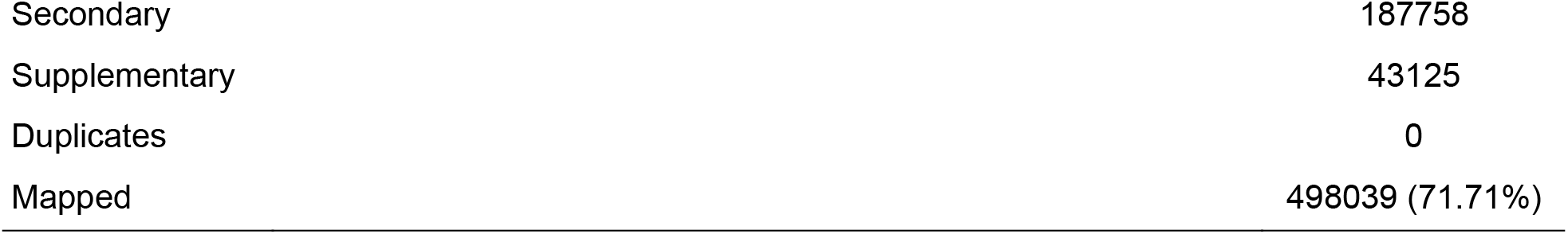
Summary statistics for the de novo transcriptome used as full-length evidence.

**Table S3.** Repeat content.

|  |  | **Total elements** | **Element length (bp)** | **Coverage (%)** |
| --- | --- | --- | --- | --- |
| SINEs |  | 0 | 0 | 0 |
| LINEs |  | 702,573 | 578,885,792 | 2.09 |
|  | LINE1 | 370,868 | 404,306,289 | 1.46 |
| LTR |  | 7,347,127 | 11,559,544,152 | 41.77 |
|  | ERV_classII | 42,505 | 35,061,926 | 0.13 |
| DNA |  | 399,110 | 310,688,087 | 1.12 |
| Unclassified |  | 18,838,504 | 8,937,651,555 | 32.295 |
| Total interspersed |  |  | 21,386,769,586 | 77.28 |
| Small RNA |  | 45,706 | 40,193,439 | 0.145 |
| Simple repeats |  | 44,012 | 46,361,512 | 0.17 |
| Total |  |  | 21,473,324,537 | 77.59 |

**Table S4.** Genome annotation approaches.

| Run | Total transcripts | Total genes | Gene N50 (bp) | BUSCO (run in protein mode) | EnTAP (%) | Mono: multi | Max intron (Mb) | average exons per gene |
| --- | --- | --- | --- | --- | --- | --- | --- | --- |
| StringTie2 (SR) | 63,123 | 51,844 | 2,281 | C:70.9% [S:43.3%, D:27.6%], F:15.2%, M:13.9% | - | 0.34 | 1.39 | 4.7 |
| StringTie2 (Hybrid) | 62,936 | 47,132 | 1,807 | C:71.4% [S:37.4%, D:34.0%], F:16.5%, M:12.1% | - | 0.33 | 1.02 | 4.2 |
| BRAKER (SR) | 636,628 | 631,708 | 963 | C:45.0% [S:35.6%, D:9.4%], F:30.8%, M:24.2% | - | 1.03 | 0.14 | 2 |
| BRAKER (SR)<br>eggNOG Filtered | 219,474 | 216,079 | 1,164 | C:45.0% [S:35.7%, D:9.3%], F:30.9%, M:24.1% | - | 0.70 | 0.14 | 2.7 |
| StringTie2 (SR)<br>eggNOG +<br>Transdecoder<br>filtered | 48,567 | 27,336 | 1,578 | C:70.5% [S:43.2%, D:27.3%], F:15.1%, M:14.4% | 0.85 | 0.22 | 1.39 | 5.9 |
| StringTie2 (Hybrid)<br>eggNOG +<br>Transdecoder<br>filtered | 45,380 | 31,953 | 1,515 | C:70.3% [S:45.6%, D:24.7%], F:16.0%, M:13.7% | 0.81 | 0.23 | 1.02 | 5.3 |
| Final annotation | 47,911 | 27,010 | 1,578 | C:70.6% [S:43.2%, D:27.4%], F:15.1%, M:14.3% | 0.87 | 0.22 | 1.39 | 6 |

**Table S5.** Summary statistics for NLR identification.

|  | Only NBARC | Missing N-terminal | Missing LRR | Complete | Total |
| --- | --- | --- | --- | --- | --- |
| Denovo-assembled transcriptome | 27 | 24 | 14 | 24 | 89 |
| Direct genome (NLR-Annotator) | 616 | 771 | 257 | 595 | 2239 |
| Genome annotation | 56 | 57 | 37 | 88 | 238 |
| BRAKER gene model | 0 | 0 | 0 | 27 | 27 |

